# Molecular and ultrastructural characterization of the intramuscular nerve cords of the octopus arm

**DOI:** 10.64898/2026.06.06.730610

**Authors:** John Benedict, Mox Engelman, Meaghan Klos, Robyn J. Crook, Gabrielle C. Winters-Bostwick

## Abstract

Cephalopod arms are controlled by a distributed peripheral nervous system comprising the axial nerve cord (ANC), subacetabular ganglia associated with each sucker, four longitudinal intramuscular nerve cords (INCs) embedded within the arm musculature and oblique connectives (OCs) running between INCs. Despite their prominent anatomical position and proposed roles in local sensorimotor integration and inter-arm coordination, the INCs remain poorly characterized with respect to cell-type composition and molecular identity. Here, we report the first integrative characterization of INC structure and composition in *Octopus bocki* by combining serial block-face scanning electron microscopy (SBEM) with multiplexed hybridization chain reaction (HCR) *in situ* labeling. We show that oral and aboral INCs share a consistent internal organization comprising distinct cell body regions, a peripheral tract layer, and a central synaptic zone. Both oral and aboral cords contain morphologically diverse cell populations, including abundant bipolar neurons with long unbranched processes and a second class of neurons with highly branched processes bearing bouton-like enlargements. On the molecular level, the sampled INCs are enriched for glutamatergic and buccalin-positive cells, and express abundant glia-associated transcripts. In contrast to the ANC, cholinergic, dopaminergic, serotonergic, and octopaminergic markers were not detected above background. We also characterize the relationship between the INCs and adjacent oblique connectives (OCs), showing that these structures run in close proximity but remain physically separate within the sampled high-resolution volume, with no shared fibers or crossing processes detected across the observed boundary. Together, these data establish a cellular and molecular framework for the INCs and clarify their relationship to neighboring peripheral pathways.

## Introduction

The complex architecture of nervous systems in octopus arms supports sophisticated motor control and rich somatosensation distributed across peripheral circuits and central brain pathways. Each arm contains extensive neural circuitry embedded within a muscular hydrostat that permits near infinite degrees of freedom, enabling flexible behaviors including reaching, object manipulation, exploration, and defensive movements^1–8^. Classic and modern work establishes that many sophisticated arm behaviors can be generated or shaped by local sensorimotor loops, with the brain providing commands and modulatory control rather than specifying every detail of limb kinematics^9–12^. Recent neuroanatomical, molecular, and physiological studies have renewed interest in identifying the specific pathways and cell types that support this distributed control, including the axial nerve cord (ANC), subacetabular ganglia, and associated peripheral pathways ^13–15^. However, the intramuscular nerve cords (INCs) - longitudinal nerve tracts embedded deep within the arm musculature - remain comparatively poorly resolved beyond gross anatomical descriptions^16–19^ and recent discoveries of long-range inter-arm continuity^20^, limiting mechanistic models of how sensory and motor information is integrated within and across arms.

Classical anatomical descriptions place INCs as major longitudinal elements of the peripheral arm nervous system and associate them with proprioception and local motor coordination^16,21^. Their deep intramuscular position is well suited to influence local motor recruitment, coordinate reflex-like signaling, and distribute sensorimotor information along the arm. More recent work in *Octopus bimaculoides* has suggested that INC pathways may also contribute to inter-arm coordination through peripheral routes that are independent of canonical central pathways^20^. That and subsequent studies^22^ confirm that oral INCs extended proximally beyond the arm base and are continuous with corresponding pathways in arms two positions away, revealing a stereotyped peripheral architecture capable of linking distant arms across the crown without involving the central brain or ANCs^20^.

Ultrastructural analyses of the octopus arm have also found that the peripheral nervous system is organized by repeating architectural motifs rather than isolated tracts. Multiple longitudinal pathways coexist in close proximity, including the ANC, intramuscular cords, sucker-associated nerves, and oblique connectives (OCs) that travel through the arm musculature^13^. This organization implies that the INCs are not simply accessory bundles embedded in muscle but components of a broader peripheral network with the potential to coordinate local reflexes, influence the recruitment of muscle groups, and distribute sensorimotor signals through the musculature.

Despite their anatomical prominence and strategic position, the INCs have received only limited attention regarding cell-type composition or transmitter phenotype, including immunohistochemical identification of pChAT-positive neurons^23^. Addressing this gap requires approaches that preserve spatial context while resolving cell identity. Over the past decade, cephalopod neuroscience has expanded rapidly through transcriptomics, single-cell approaches, and multiplexed *in situ* labeling, which are beginning to yield cell-type maps and candidate transmitter phenotypes in central and peripheral nervous tissues^14,23–30^. In parallel, advances in volume electron microscopy have enabled detailed reconstruction of cephalopod neural microanatomy in the central brain, in particular the connectome of the Octopus vulgaris vertical lobe, illustrating novel interneuron types, synaptic motifs, and the organization of a learning and memory network at nanoscale resolution^31,32^ and in the arms^13^, clarifying how axonal tracts, neuropil domains, glia-like elements, and connective tissue are organized in relation to the arm musculature. However, molecular identity and ultrastructural organization have rarely been combined within the same peripheral structures. This gap is especially pronounced for the INCs, where the absence of molecularly anchored cell-type information has limited interpretation of their functional role and their relationship to neighboring arm circuits.

In the present study, we characterize the octopus INCs by combining molecular cell-type identification with ultrastructural analysis in the miniature tropical octopus *O. bocki*. The very small adult size of this species (mantle length ∼1cm) allows us to examine mature neuroanatomy on a scale smaller than that of hatchlings of many commonly studied larger species. We use SBEM to define INC microanatomy, reconstruct local tract organization, and identify distinct cellular morphologies within oral and aboral cords. In parallel, we use multiplexed HCR in situ labeling on fixed tissue sections to assign molecular identities to major neuronal populations using markers associated with neurotransmitter and neuropeptide signaling. This combined approach allows us to link cellular morphology and tract architecture to molecularly defined neuronal classes, compare INC composition between the oral and aboral cords, and begin to resolve how these pathways relate to neighboring peripheral structures. By examining regions where OCs and INCs run in close proximity, we also assess whether these pathways are structurally integrated or remain anatomically distinct. Together, these data establish a cellular and anatomical framework for one of the least characterized major pathways in the octopus arm nervous system.

## Materials and Methods

### Animal acquisition, husbandry, and euthanasia

Animal acquisition, husbandry, and euthanasia followed previously published protocols for *O. bocki* arm tissue studies^13,14^. Adult *O. bocki* (Figure 1A) of undetermined sex were obtained from SeaDwelling Creatures (Los Angeles, CA, USA) and housed individually in enriched enclosures containing substrate, shells, coral rubble, and artificial plants. Animals were fed 3 to 4 live grass shrimp daily. Individuals were monitored routinely for signs of poor health or compromised welfare and were maintained in the laboratory for at least one week before tissue collection. All animals selected for tissue collection were feeding normally and showed no obvious signs of senescence or declining health.

**Figure 1.**
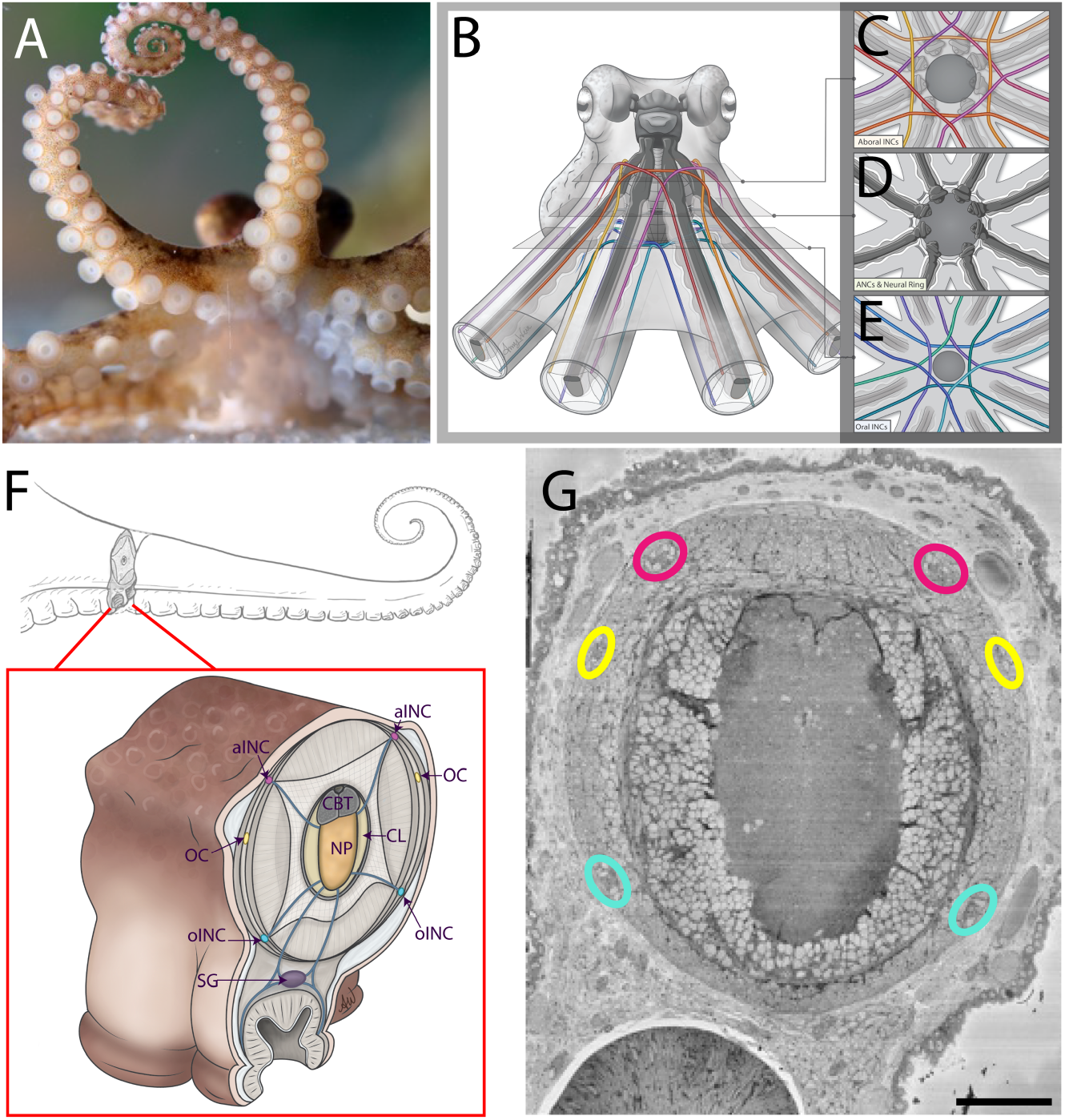
INCs and OCs run peripheral to the ANC in octopus arms. A. *Octopus bocki* arms showing typical two-row arrangement of suckers. B-E- Schematic illustrating the 3D spatial distribution of the oral and aboral INCs in arms and as they converge and extend to other arms. Panel B orients the oral (cool tones) and aboral (warm tones) INC organization in the context of whole animal including the ANCs as they bypass the buccal mass and extend down the arms. Panels C-E are rotated to a top down cross section of the central organization of the aboral INCs (C), ANCs (D), and oral INCs (E). F. Schematic illustration of a cross section of an octopus arm with further magnification depicting major neuroanatomical structures. G. A single section from an overview stack of the whole arm, showing the main axial ganglion and the four smaller intramuscular nerve cords; the two aboral cords in magenta, oral cords in cyan, and two oblique connectives in yellow. Abbreviations: NP, neuropil of the ANC ganglion; CL, cortex layer of the ANC ganglion; CBT, cerebrobrachial tracts; SG, subacetabular ganglion; oINC, oral intramuscular nerve cord; aINC, aboral intramuscular nerve cord; OC, oblique connective.

For tissue collection, animals were deeply anesthetized by immersion in isotonic magnesium chloride solution (330 mM MgCl2.6H2O in reverse osmosis deionized water). For EM preparations, the solution additionally contained 1% (v/v) ethanol. To minimize disturbance, animals were transferred into the anesthesia bath while remaining inside the shell or PVC shelter used as a den. Fifteen minutes after respiration ceased, animals were killed by decerebration.

Ethical note: Octopuses are invertebrates and are therefore excluded from regulatory oversight in the USA; no IACUC protocol was required for this study. However, all procedures adhered to Directive 2010/63/EU and ARRIVE guidelines for characterizing standards of care, humane endpoints, and experimental procedures.

### Electron microscopy datasets

Electron microscopy datasets analyzed here were generated from *O. bocki* arm tissue using serial block-face scanning electron microscopy. Tissue preparation, image acquisition, and 3D reconstruction followed the protocol described in Neacsu and Crook^13^. After animals were decerebrated, their two front arms (L1 and R1) were removed, and small arm segments approximately 1 mm in length were fixed immediately in 4% paraformaldehyde plus 2.5% glutaraldehyde in 0.1 M cacodylate buffer, pH 7.2, for 5 to 7 days at 4 °C. Fixed samples were shipped on ice to the 3DEM Ultrastructural Imaging and Computation Core at the Cleveland Clinic Lerner Research Institute (Cleveland, OH, USA), where tissue was post-fixed with osmium tetroxide, dehydrated through a graded ethanol series, stained *en bloc* with uranyl acetate, and embedded in epoxy resin.

Samples were sectioned and imaged in cross section using either a Thermo Fisher Scientific Teneo VolumeScope or a Zeiss Sigma VP microscope equipped with a Gatan 3View in-chamber ultramicrotome. Two datasets were acquired, one centered on the left oral INC, and one on the left aboral INC. Both datasets were acquired from the same distal region of the L1 front arm. The oral INC dataset comprised 761 serial sections imaged at 7,7,70nm xyz, dimensions 89,131,53 μm (https://webknossos.org/links/7uAmVu-za0VJ5Pnj). This volume captured a passing oblique connective as it descended orally, made contact with the oral INC and passed beyond it along a centro-oral trajectory. The aboral INC dataset comprised 795 serial sections imaged at 7,7,70nm xyz, dimensions 141,134,55 μm (https://webknossos.org/links/3GW4meFvrXr1_rjk).

Image registration, segmentation, and volumetric visualization were performed in webKnossos^33^. Sections were aligned using the AI-assisted alignment workflow, and structures were annotated using automated tools for segmentation by all authors, and each segment was cross-checked and corrected manually if needed. Surface renderings of approximately 230 individual cell structures were generated then categorized based on morphology. Final volumes and overlays were exported to Amira 10.1 for visualization.

### Tissue preparation for HCR

Tissue preparation for HCR followed the protocol described in Winters-Bostwick et al^14^. After animals were decerebrated, their arm tissues were dissected and fixed overnight at 4 °C in 4% paraformaldehyde in PBS. Samples were rinsed in PBS, washed in PTW (1x PBS, 0.1% Tween-20), dehydrated through a PTW:methanol series, and stored at -20 °C. Before hybridization, tissues were rehydrated through the reverse series, rinsed in PTW, post-fixed for 1 h in 4% paraformaldehyde in PBS at 4 °C, cryoprotected in 30% sucrose in PBS, embedded in Tissue-Tek O.C.T. compound (Sakura Finetek, cat. no. 4583), frozen, and sectioned at 50 μm. Serial sections were collected on slides, dried, circled with an ImmEdge hydrophobic barrier pen (Vector Laboratories, cat. no. H-4000), and processed in a humidified chamber. Sections were washed in PTW for 10 min, permeabilized for 10 min in 0.3% Triton X-100 in PBS, and washed again in PTW before hybridization.

### Hybridization chain reaction in situ labeling

HCR RNA *in situ* hybridization was performed on slide-mounted sections using the Molecular Instruments protocol for generic samples on a slide (rev. 9, 2023-02-13), following the HCR v3.0 framework^34^ and as previously described in Winters-Bostwick et al al^14^. Custom probe sets were designed through Molecular Instruments using transcript sequences identified in that prior study. To improve signal in octopus arm tissue, probes were used at 16 nM. Salmon sperm DNA was added to the amplification buffer at 100 μg/mL during pre-amplification and amplification steps, and the pre-amplification incubation was extended from 30 min to 1 h to reduce nonspecific background. Remaining hybridization, wash, and amplification steps followed the manufacturer’s instructions. Sections were mounted in Fluoromount-G with DAPI (Thermo Fisher, cat. no. 00-4959-52). We re-imaged INCs from existing sections collected for a prior study^14^ and quantified presence, absence, and relative abundance of cell types by transcript across structures.

### Confocal imaging

Fluorescent images were acquired on a Leica Stellaris 5 confocal microscope on a DMi8 (supported by NSF MRI 2018239) using an HC PL APO CS2 40x/1.30 oil objective and LAS X software (v4.6.0.27096). DAPI was imaged with a 405 nm diode laser; other fluorophores were acquired using the white light laser. Images were collected at 1024 × 1024 resolution with a pixel dwell time of 0.6 μs using bidirectional scanning. Image processing was performed in FIJI^35^.

## Results

### Overall organization of the intramuscular nerve cords

Overview EM datasets confirm that *O. bocki* arm cross sections contain four intramuscular nerve cords arranged in two bilateral pairs: two oral cords positioned on the ventral (toward the sucker) face of the arm and two aboral cords positioned dorsally (Figure 1B-F). This arrangement is consistent with prior anatomical descriptions of INC position in octopus arms^16,20,21,36^. At low resolution, each INC appears as a compact, elongated structure embedded within the oblique musculature, clearly distinguished from surrounding muscle and connective tissue by the density of nuclei within it (Figure 1G, 2A).

High-resolution SBEM datasets (7 × 7 × 70 nm voxel size) from oral and aboral INCs revealed that each cord shares a consistent internal organization (Figure 2 B,C). Four spatially distinct zones are visible in cross section: two cell body zones positioned on the oral and aboral margins of the cord; a tract layer on the peripheral lateral face, containing tightly bundled, longitudinally running processes; and a synaptic zone on the central face, closest to the ANC, containing dense, fine, vesicle-filled processes (Figure 2D). This general organization was observed in both the oral and aboral INCs.

**Figure 2.**
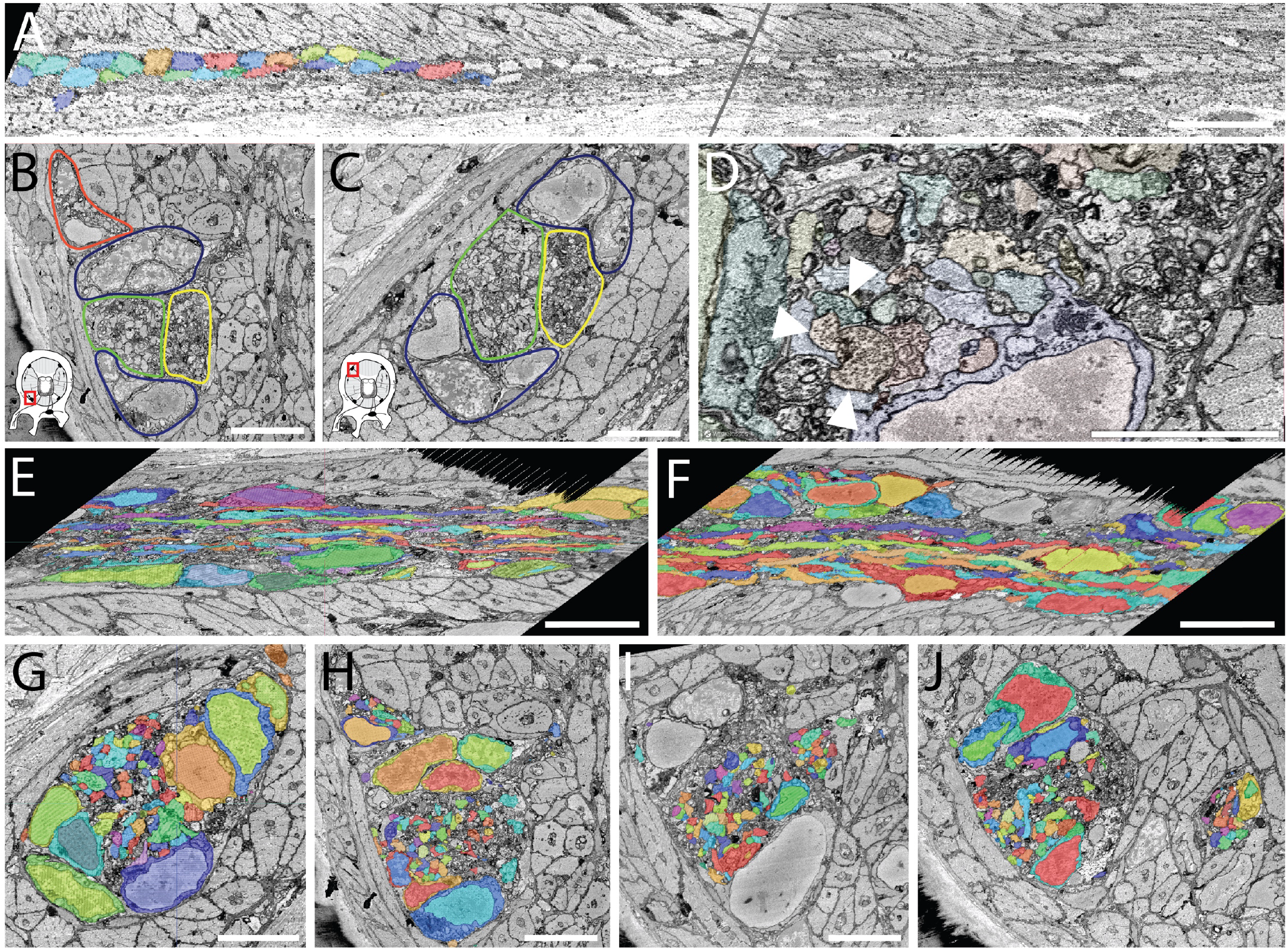
Internal organization of the intramuscular nerve cords. A. Serial blockface EM of the oral INC at low resolution, showing cell bodies (some colored) evenly spaced along the cord. B. High-resolution imaging of the oral INC (7×7×70nm) reveals INCs have consistent organization. In this view an oblique connective (red) is present on the aboral margin. There are two cell body zones on the oral and aboral margins (blue), a tract region on the peripheral lateral side (green), and a synaptic zone containing fine, vesicle-filled processes on the medial side, closest to the main axial nerve cord (yellow). C. The same organization is present in the aboral INC. D. A magnified view of the synaptic zone, showing putative synaptic contacts between vesicular boutons and other processes E. Longitudinal view of the segmentation of the aboral cord. F. Longitudinal view of the oral INC, showing segmentation of cell bodies and processes. G. Cross-section of the aboral INC, showing AI-assisted manual segmentation. H-J. The oral INC, with an oblique connective bypassing to the central, oral aspect. Scale bars: A, 150 μm; B-J (except D), 20 μm. D, 5 μm

Longitudinal sections confirmed that the tract and synaptic zones extend along the cord without obvious interruption (Figure 2E,F). The aboral cord in the volume sampled (Figure 2G) was not in a region close to an oblique connective, whereas the oral cord was imaged just as a descending OC was making contact with the INC (Figure 2H) and the OC continued alongside the INC (Figure 2I) before separating on the medial oral margin of the INC and traveling toward the arm midline in the muscle (Figure 2J). These data establish that the INCs are not simple nerve bundles but organized structures with reproducible internal compartmentalization, nor do the OCs provide simple linking tracts between oral and aboral INCs.

### Neuronal morphology in the INCs

Manual segmentation and AI-assisted surface rendering of cells within the high-resolution INC volumes allowed us to segment and reconstruct 132 individual cells or cell compartments across oral and aboral cords (Figure 3), focusing primarily on large structures and somata. Based on soma morphology, nuclear characteristics, and process architecture, most neurons fell into two broad morphological classes, with additional less common types also observed.

**Figure 3.**
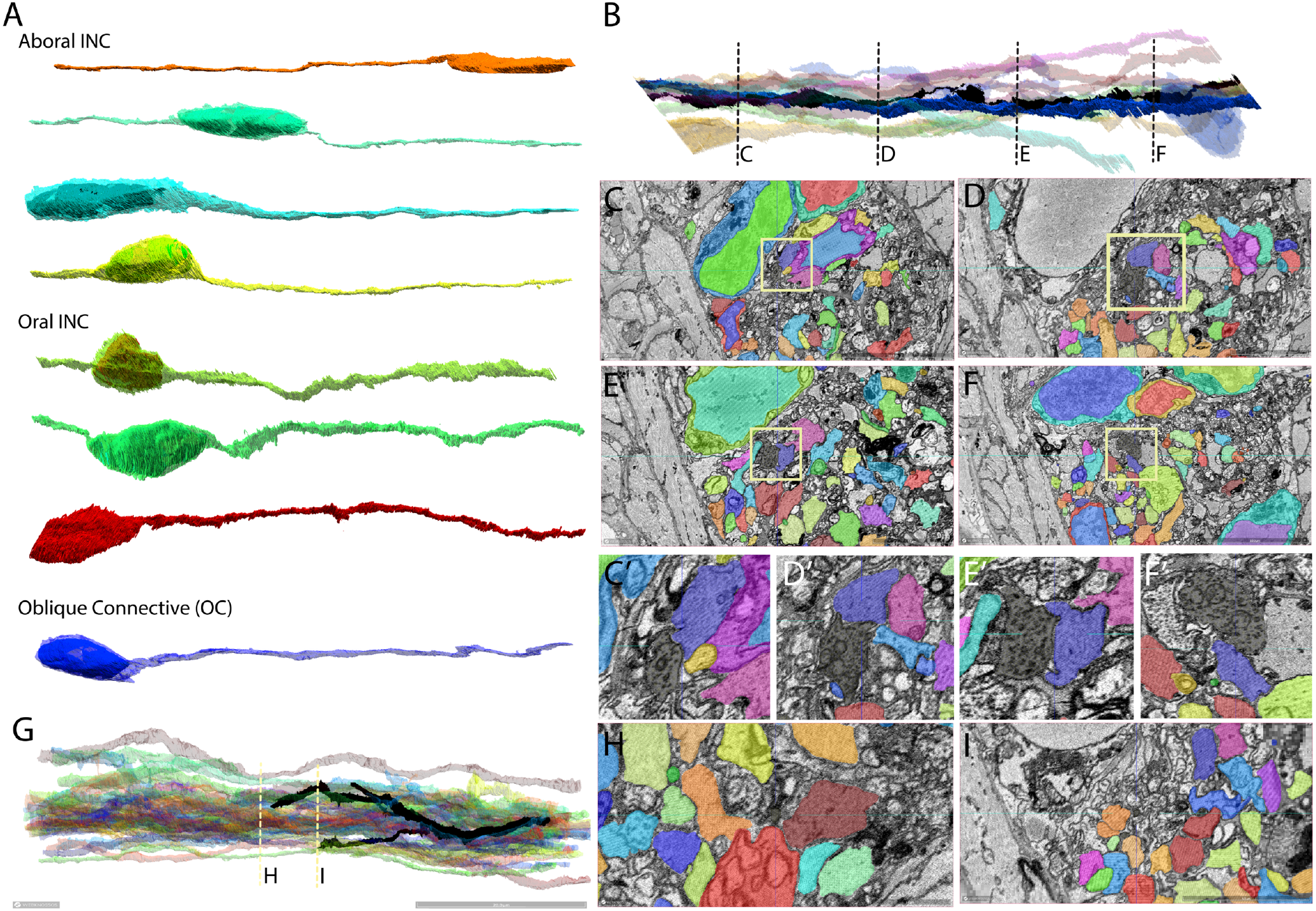
Reconstructions of cell architectures from 3DEM. A. Many cells were simple bipolar cells with long, unbranching processes running in the tract zone of each cord, with similar gross morphological characters in all three cords. B. Segmentation of processes running in the tract zone show that the majority of structures are likely bipolar cell processes. Although the tract and plexiform zone remain largely separate, there are occasional varicose processes of quite different morphology, which are dense with synaptic vesicles. The two opaque structures show a typical bipolar cell process (blue) and a vesicular process (black) in close contact over the length of the tissue block. Transparent meshes are other bipolar processes making single contacts with the same vesicular process). Dotted lines show the position of the xy images in C-F. C’-F’ are close-up views (boxes), showing the close contact between blue and black processes over ∼700 sections. G. The majority of simple bipolar processes extend out the end of the block in both directions without branching. Only a few processes terminated in the block, often making one or two branches and terminating among vesicular processes traveling within the tract zone. Dotted lines show the position of xy images in H and I. The termination of the black process is centered under the cross hairs, where it is engulfed in vesicle-dense varicosities from other, untraced structures.

The most abundant class comprised simple bipolar neurons with large nuclei and two long, unbranched processes that run within the tract layer of the cord, extending through the full sampled volume without obvious arborization (Figure 3A). These cells were present both in oral and aboral cords. Their process morphology is consistent with long-range relay or projection neurons, although definitive evidence of synaptic output from these cells was not obtained within the sampled volume. Simple bipolar cell processes comprise the majority of the tract layer, and these processes rarely deviated into the plexiform zone on the medial side of the INC, where presumably they would be in contact with the dense vesicular processes that comprise the majority of projections in this region. However, processes that are dense with vesicles are present at low abundance in the tract zone (Figure 3B), where they make repeated, close contacts with a small subset of bipolar cell processes. The most common pattern of contacts we observed was a close pairing between a single vesicular process and one bipolar process (Figure 3C-F), with other bipolar processes making single contacts along the length of the varicose process. Only a few bipolar cell processes appeared to terminate in the block (Figure 3G), typically making one or two short branches before terminating in the tract zone without forming obvious boutons with vesicles. Terminations were generally among untraced, small varicosities that were dense with synaptic vesicles (Figure 3H, I) suggesting these endings are postsynaptic.

We also found a second, less abundant class of neurons, still generally bipolar in morphology but with more complex process architectures in all three cords (Figure 4A). In some cells there were multiple short, unbranched dendrites emanating from the cell body, and in others of roughly the same morphology there were no cell body dendrites, potentially suggesting a subdivision within complex bipolars. Both classes have thick primary dendrites, which were characterized by multiple bouton-like enlargements and sometimes branched offshoots (Figure 4B), which ran in both directions along the cord. These processes were often filled with mitochondria and large, clear vesicles (Figure 4C-F).

**Figure 4.**
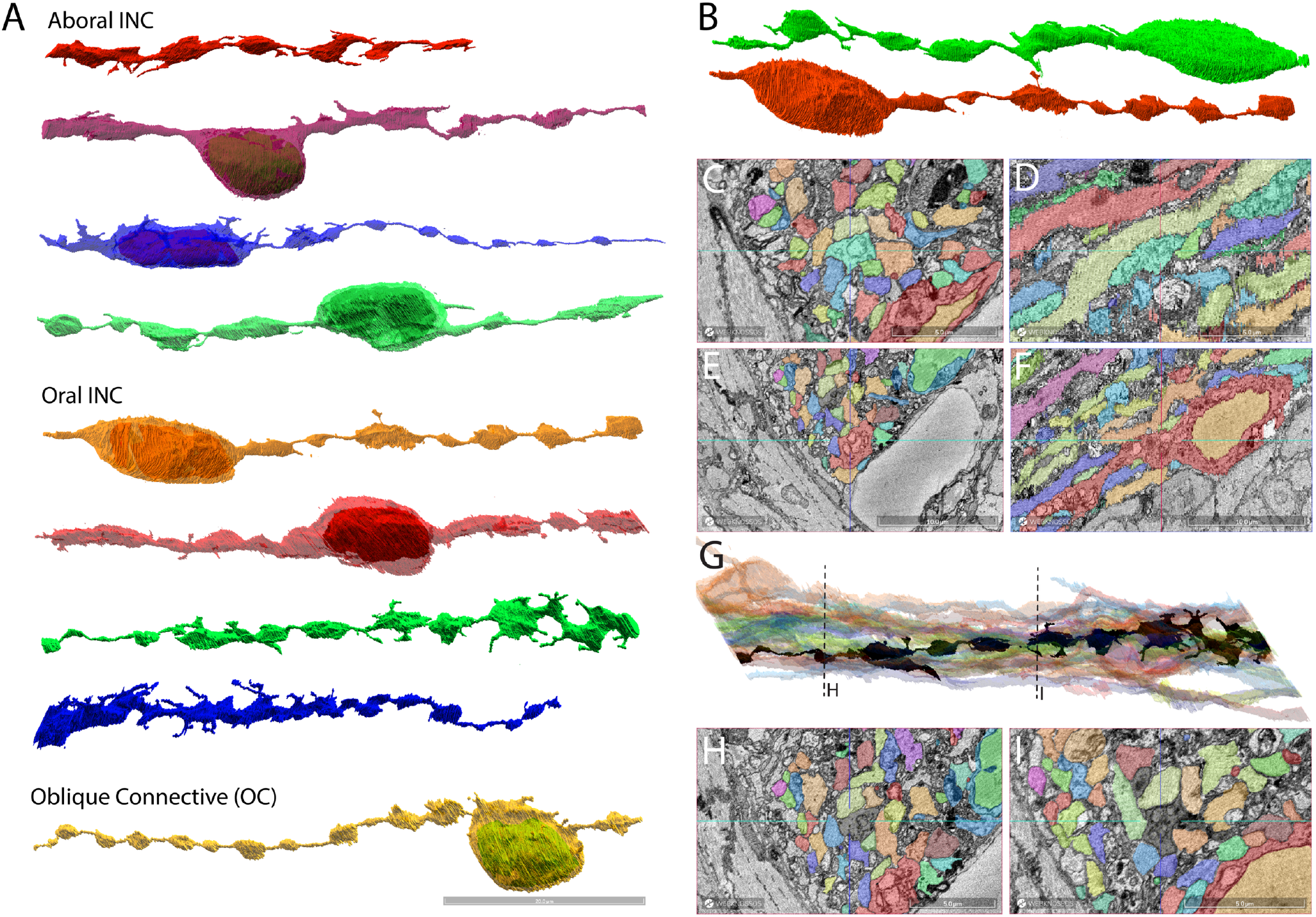
Reconstructions of complex cell architectures from 3DEM. A. Cell segments from each of the three cords, showing similar morphologically-complex bipolar cells in all three. Some differences among them are apparent in the degree of dendritic branching from the soma and from varicosities along each extending process, suggesting additional division within the broad class we term complex bipolars. B. An example of two cells traced from the oral INC, running in opposite directions in the cord. The dendrite is thick as it departs from the soma and then forms varicosities along its length. The axon is noticeably thinner, a feature also described by Graziadei in branching cells in the INC. C. An xy view of the green cell in B, centered under the cross hairs and showing the convoluted shape of the dendrite and its large mitochondria. D. The same cell in yz orientation, revealing mitochondria rich, large boutons along the length of the presumptive dendrite. No synaptic vesicles are apparent in the process. E. An xy view of the red cell in B, showing highly similar ultrastructural features to the green cell, suggesting these two cells are of the same type. F. A yz view showing the thick dendritic trunk as it departs the soma, and the same internal feature of many large mitochondria. G. One of the highly dendritic orphan processes (no cell body in the block) shown in green in A, here shown in opaque black for visibility. Surrounding it in transparent colors are all the cells with which it makes physical contact through the block (39 in total). Dotted lines show the position of the xy view in H and I. Notably, the majority of close contacts of either the main dendrite or terminal contacts of the short branches are onto other clear processes, not vesicular boutons. The function of these complex bipolar cells remains unclear.

Most varicosities of these within the sampled volume were clear, lacking obvious synaptic vesicles, suggesting that they may represent postsynaptic contacts, although they were confined primarily to the tract zone rather than the plexiform zone of the cord. Highly branched processes occurred in about half of all complex bipolars, and these cells made physical contact with between 30 and 40 other traced structures (which often excluded small vesicular processes due to their very small size and difficulty of following through serial sections, Figure 4G). We found multiple examples of the fine branching dendrites of these cells terminating among other clear processes presumably belonging to simple bipolars (Figure 4H,I) but the function of these contacts remains obscure.

Fine, vesicle-filled processes in the typical en-passant configuration were abundant in the synaptic zone, but were difficult to trace through the full block with confidence due to their extremely small diameter in between varicosities (many less than 100nm). Others were orphan fibers projecting into the cord from outside the sampled block, potentially originating from cells in the ANC or other neighboring pathways. Typical bipolars, complex bipolars and vesicular processes were identified in the oral and aboral INCs and in the oblique connective, suggesting shared cellular features across these peripheral structures. Additional cell types with unusual nuclear staining patterns or morphologies distinct from either major class were also observed in the oral and aboral INCs, but were less numerous; these are described further in the context of glial cell characterization below.

The synaptic zone, positioned on the central face of the cord nearest the ANC, contained dense accumulations of vesicle-filled processes forming putative synaptic contacts with other processes (see Figure 2D); a small minority could be traced through the block but the majority entered through a nerve bundle projecting from the ANC. Notably, no cell bodies within the sampled INC volume sent processes back toward the ANC. This directional asymmetry-vesicular input arriving from the ANC side with no identified reciprocal projection- is consistent with a model in which the INC receives descending or collateral input from the ANC rather than providing a direct ascending projection, though the limited tissue volume sampled prevents a definitive conclusion.

### Neuronal molecular identity in the INCs

To complement the morphological data, we used multiplexed HCR labeling to characterize the neurotransmitter and neuropeptide phenotypes of INC neurons. We applied probe sets for vesicular glutamate transporter (vGlut), vesicular GABA transporter (vGAT), dopaminergic, octopaminergic, serotonergic, and several subtypes of peptidergic cells, including bradykinin-like peptide, FLRIamide, buccalin, and FCAP (Feeding Circuit Associated Peptide), and compared relative abundance across the oral INC, aboral INC, oblique connective, and ANC (Table 1).

**Table 1.**
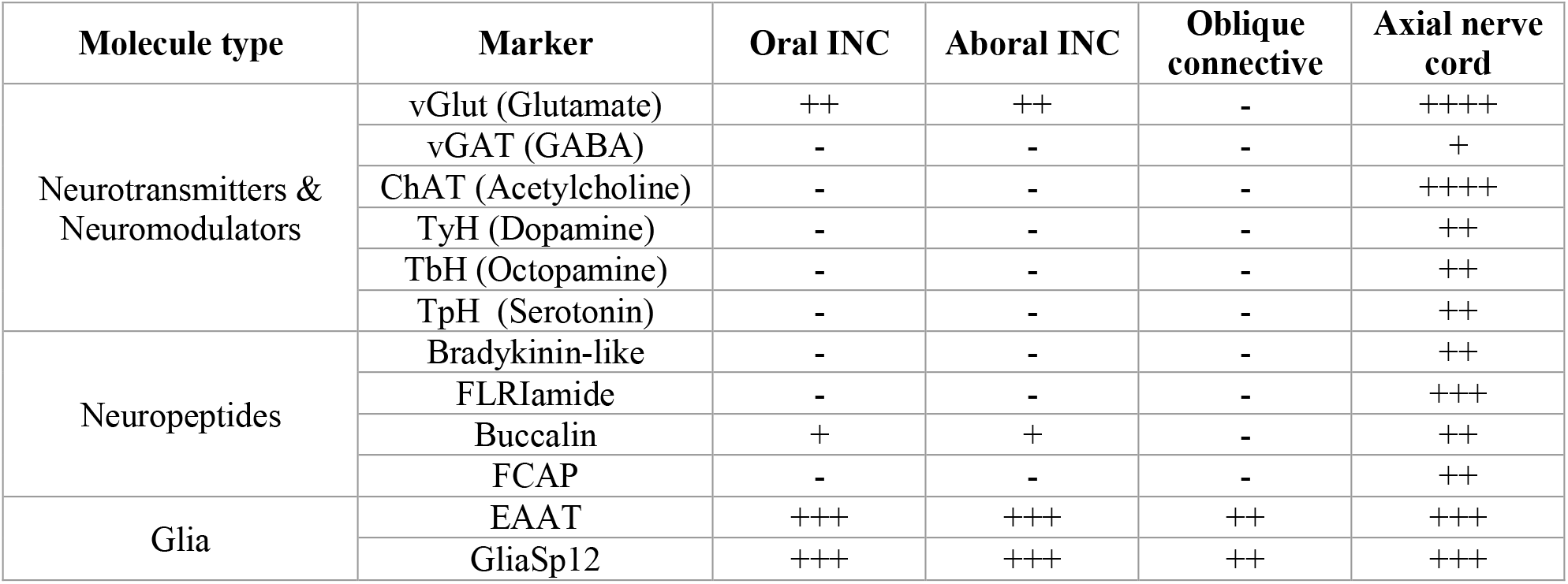
Relative abundance of neurotransmitter, neuromodulator, neuropeptide, and glial transcripts in oral INC, aboral INC, oblique connective, and axial nerve cord of *Octopus bocki*. Scores are semi-quantitative estimates from the sampled material and should not be interpreted as precise cell counts.

The oral and aboral INCs showed broadly similar molecular profiles. vGlut-positive cells were present in both cords at moderate abundance (Figure 5A-C), consistent with a glutamatergic neuronal population being a significant feature of the INCs. Of the monoamine markers tested, dopamine, octopamine, and serotonin did not produce labeling above background in either sampled INC, in contrast to their well supported presence in the ANC^14^.

**Figure 5.**
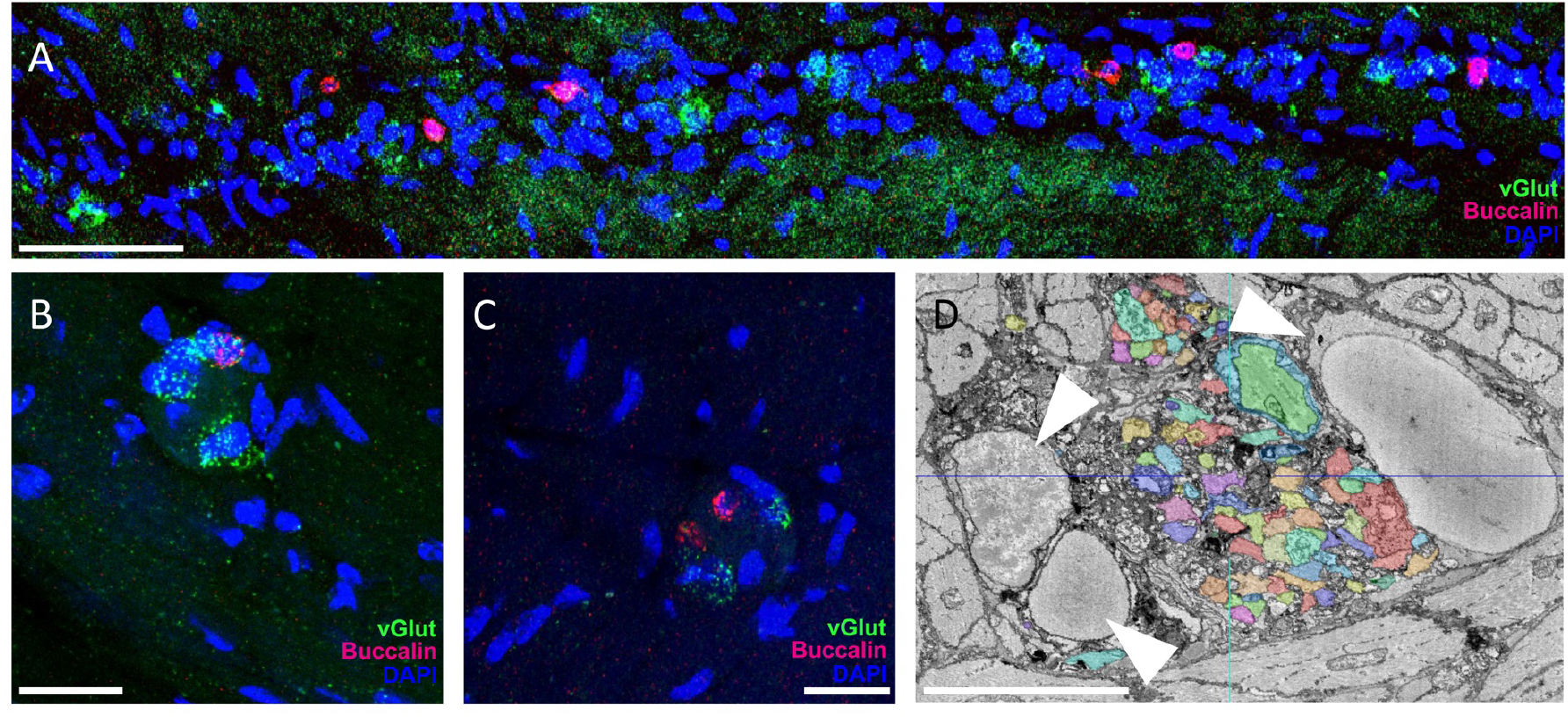
Neuronal diversity in the INCs. Glutamatergic (vGlut, green) and peptidergic (Buccalin, red) markers were detected in cell bodies throughout the INCs of both oral (A: longitudinal; B: cross section) and aboral (C) cords. Molecular cell-type identity in the INC parallels the morphological diversity seen in EM data (D, white arrows). Scale bars: A-10 μm; B-D 20 μm.

ChAT, which labels acetylcholinergic cells, was expressed abundantly in the ANC in the same sections in which we did not detect ChAT signal in the INCs (Figure S1). This contrasts with prior immunohistochemical work in *O. vulgaris* showing ChAT-immunoreactive structures in the INC ^23^. Among neuropeptides, buccalin-positive cells were identified in both oral (Figure 5B) and aboral (Figure 5C) INCs at low abundance, while bradykinin-like, FLRIamide, and FCAP labeling was absent or below detectable levels. In the sampled sections, buccalin-positive and vGlut-positive cells appeared as non-overlapping populations: buccalin-positive cells did not show detectable vGlut signal, and vGlut-positive cells did not express detectable buccalin (Figure 5B-C). Even this partial molecular survey suggests that the INC houses more cellular diversity than has previously been appreciated, paralleling the range of soma morphologies seen in the EM data (Figure 5D).

The oblique connective presented a more ambiguous molecular profile, largely due to the lower density of cell bodies and practical challenges associated with imaging structures of its size and orientation in these tissue sections. vGlut labeling was not detected above background in the OC, and other neuronal markers did not produce definitive signal.

### Glial cells are abundant in the INCs and oblique connectives

HCR labeling for two glia-associated transcripts, EAAT (excitatory amino acid transporter) and GliaSp12^28^, produced robust signal throughout both oral (Figure 6A) and aboral (Figure 6B) INCs and in the oblique connective (Figure 6A) (Table 1). Importantly, glial marker labeling was not confined to cell body zones but was distributed diffusely throughout the cord volume, including the tract and synaptic zones, in a characteristic reticulated pattern consistent with glial cell processes extending through neuropil (Figure 6A-C).

**Figure 6.**
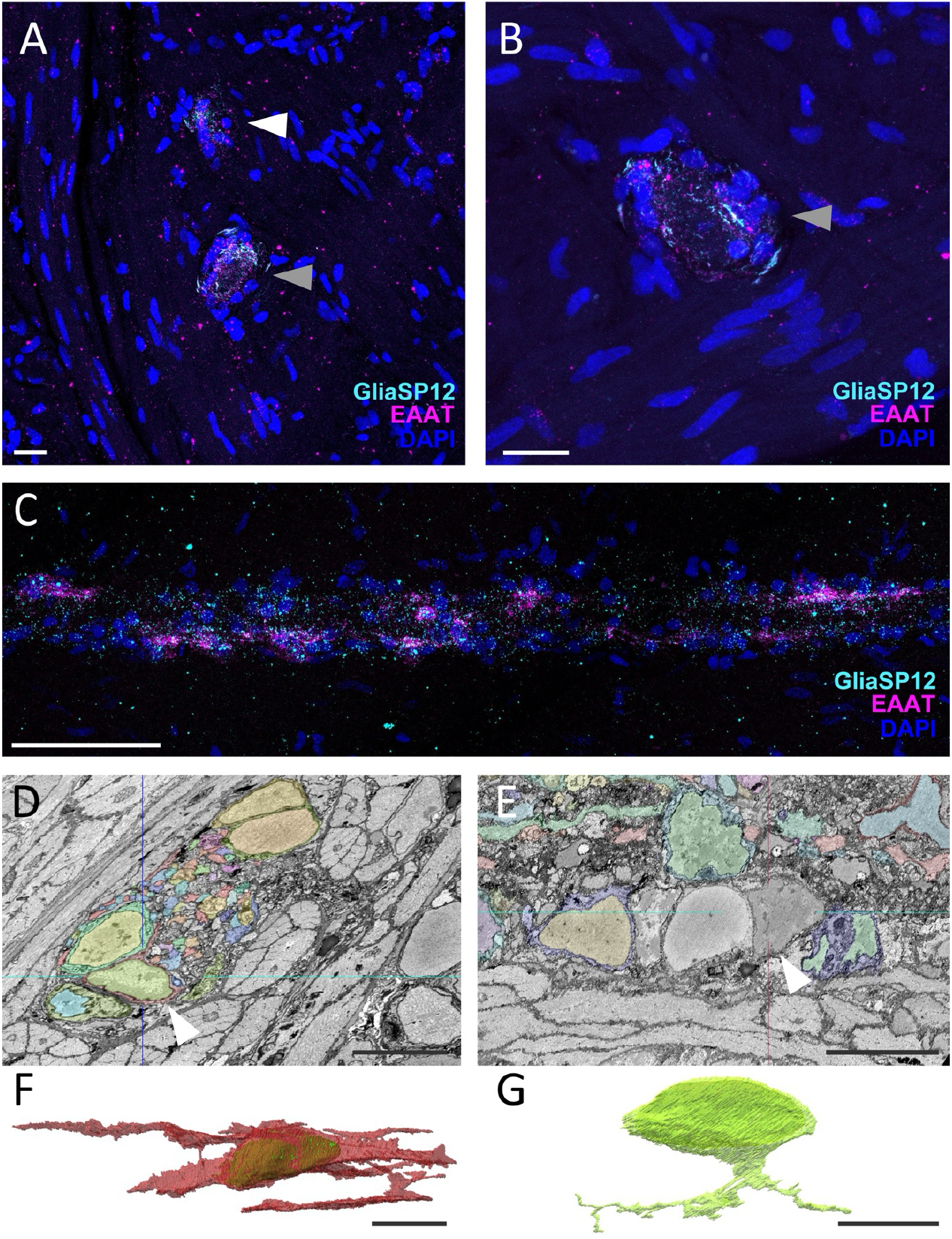
Glial cells are abundant throughout the INCs and oblique connectives. Representative cross section of an INC (gray arrows) and OC (white arrow) expressing transcripts encoding markers for glia-like cells (A-oral INC and OC; B-aboral INC). A longitudinal section of an oral INC (C) exhibits diffuse labeling throughout the INC unrestricted to neuronal somata, characteristic of glial cells. In the EM datasets we found multiple, putative glial cells characterized by long, flattened extensions that branch extensively through the volume (D white arrows & F) and wrap around the outer edge of the INCs, and other cells with shorter, branching processes or unusually dark nuclei (E white arrows & G). These cells appear similar to the protoplasmic glial cells described in other studies. Scale bars: A-B & D-G- 10 μm; C 100 μm

In the EM datasets, multiple cell types with features consistent with glial identity were identified based on ultrastructural characteristics (Figure 6D-G). One class was characterized by large, flattened cell bodies with long extensions that branch extensively through the cord volume and wrap around the outer margins of the INC (Figure 6D & 6F). This morphology is broadly consistent with protoplasmic wrapping or ensheathing glial roles described in other invertebrate nervous tissues^37–39^ and with glia-like elements reported in recent cephalopod cell-type studies ^14,25–28^. A second class showed shorter branching processes and unusually electron-dense nuclei (Figure 6E & 6G). A third, less common class was characterized by compact morphology and dark nuclear staining distinct from the two major neuronal classes. The high abundance of putative glial cells, their broad spatial distribution, and their diverse morphology collectively suggest that glial elements play a substantial structural and possibly homeostatic role in INC tissue organization.

### The oblique connective is structurally independent from the adjacent INC in the sampled volume

The high-resolution SBEM dataset of the oral INC was fortuitously acquired at a location where an oblique connective passes adjacent to the oral INC on the INC medial margin. This provided an opportunity to examine the structural relationship between the two structures at nanoscale resolution (Figure 7).

**Figure 7.**
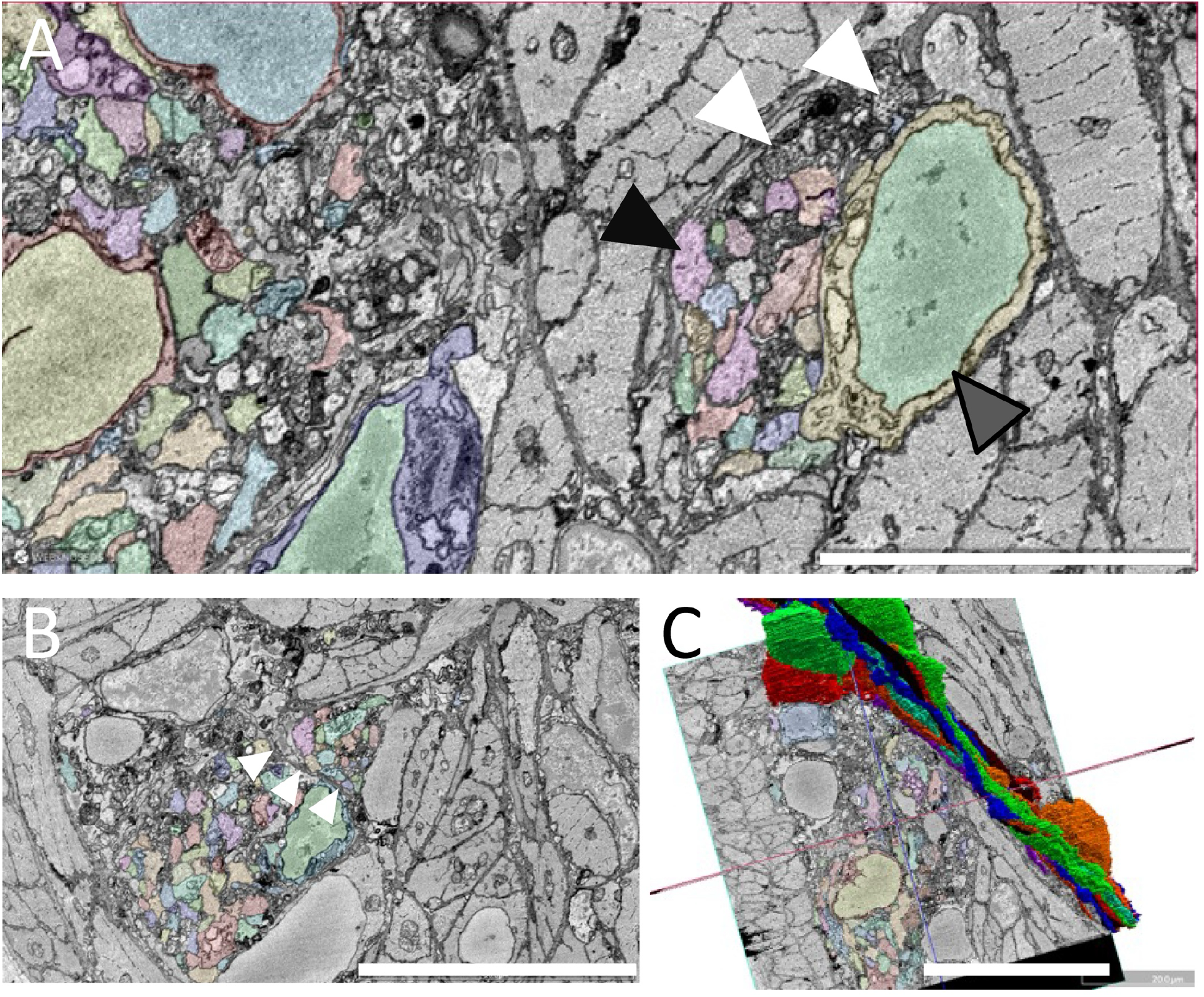
Nanoscale mapping of the oblique connective, as it passes by the oral INC. A. A view of the oral INC (left) and the OC (center). The OC contains typical tract fibers (black arrowhead) like those of bipolar cells in the INC, vesicle-filled boutons (white arrowheads) and cell bodies (grey arrowhead). B. No shared fibers travel between the two structures, and a membranous separation along the entire length in which the structures are in direct contact (white arrowhead). C. Meshes of structures traced in the OC passing by the boundary of the INC. We found no evidence of cell processes projecting from one structure to the other, although we could not rule out synaptic contact between tract processes in the OC and vesicular boutons in the INC. Scale bars: A- 100 μm; B-C 200 μm.

The OC contained tract fibers morphologically similar to bipolar cell processes observed in the INC, vesicle-filled boutons, and cell bodies belonging to typical and complex bipolar cells (Figure 7A). The OC therefore appears to be a complex structure in its own right, not simply a bundle of passing fibers. Our data show that the OC transits from the aboral margin of the oral INC where it first makes contact, along the INC medial margin in passing, and finally separates and continues into the arm musculature that is oral and medial to the INC, potentially spiraling around the oral aspect of the arm^13^. Where the two structures are in direct contact a continuous membranous boundary was visible, and no fibers were observed crossing from one structure to the other (Figure 7B). Mesh reconstructions of traced processes in the OC further confirmed that no cell processes from the OC projected into the INC, and vice versa, within the observed volume (Figure 7C). While synaptic contact between processes in the OC and vesicular boutons near the INC synaptic zone cannot be entirely excluded given the resolution limits of the dataset at the boundary, there was no evidence of shared fibers or direct structural continuity between the two pathways.

## Discussion

### INC organization reflects a consistent peripheral circuit motif

The intramuscular nerve cords of the octopus arm have long been recognized as being of potential importance to fine and gross motor control of the arms, but their cellular composition and molecular identity have remained largely uncharacterized. Here, we describe the organization of the oral and aboral INCs in *O. bocki* using complementary ultrastructural and molecular approaches, and we characterize the relationship between an oblique connective and an adjacent INC at high resolution. Our data establish that the INCs contain multiple morphologically distinct cell types, are dominated molecularly by glutamatergic neurons with smaller buccalin-positive populations, and are richly endowed with glial cells. Contrary to prior assumptions, the OC and INC remain physically separate despite running in close contact, suggesting that they represent adjacent but distinct peripheral pathways rather than a single integrated tract linking aboral and oral INCs.

The zonal organization of the INC, with cell body regions, a peripheral tract layer, and a central synaptic zone, is broadly consistent with the principle that octopus arm nervous tissues are organized into repeated, internally structured motifs rather than simple bundles^13–15,21^. The presence of a clearly defined synaptic zone on the face of the cord nearest the ANC suggests that this compartment may be specialized to receive afferent input, and the presence of vesicle-filled orphaned processes, most likely originating from neurons in the ANC, is consistent with a unidirectional organization where the INCs are a lower hierarchical level than the ANC. We found no examples of cell bodies in the INC that had processes exiting via a nerve running between the INC and ANC, but whether information also flows from the INC to the ANC remains an open question that we cannot resolve conclusively, given the spatial limits of the tissue volume sampled.

The oral and aboral INCs were highly similar in structure and molecular makeup in the two regions imaged. Despite their different positions along the dorsoventral axis of the arm, their likely exposure to different mechanical environments and the far greater density of sensory tissues on the oral side of the arm, the two cords shared the same general architecture, the same major morphological cell classes, and similar molecular profiles. This similarity suggests a primary functional role in modulating gross movement of the arm rather than finer control or coordination of suckers, or processing sensory input. Developmental or functional specializations relating to their distinct anatomical positions remain hypotheses that will require additional testing.

### Molecular phenotype of INC neurons: comparison with the ANC

The molecular profile of the sampled INCs differs substantially from that of the ANC. The ANC contains spatially stratified glutamatergic, cholinergic, dopaminergic, serotonergic, and octopaminergic populations distributed within the cortex, along with a diverse neuropeptide complement ^14^, positioning it as a major site of modulatory output to the arm. Serotonin in particular has been independently localized in arm nervous tissues by immunohistochemistry^24^, reinforcing its role as a peripheral neuromodulator in this clade. The INC’s near-absence of these markers also supports the hypothesis it sits downstream of, or alongside, this modulatory architecture rather than reproducing it. By contrast, the INCs were predominantly glutamatergic with a low-abundance buccalin-positive population.

Monoaminergic markers were not detected above background in the sampled INCs. Monoamines such as serotonin, dopamine, and octopamine are broadly associated with state-dependent modulation of motor and sensory circuits, including molluscan and other invertebrate systems ^40–42^. The lack of detectable monoaminergic marker expression in the sampled INCs suggests the INCs may function primarily in more spatially specific signaling, consistent with the prominence of glutamatergic marker expression. Glutamate is a well-established excitatory transmitter in cephalopod nervous tissues ^43,44^, and glutamatergic synapses support hippocampal-like long-term potentiation in the *Octopus vulgaris* vertical lobe^45,46^, indicating that this transmitter system can carry both fast excitatory and plastic signaling roles in octopus circuits. The dominance of vGlut in the INC, combined with the relative absence of monoaminergic markers, suggests a cord built primarily for excitatory transmission rather than broad neuromodulation, while still retaining some peptide-based modulatory capacity through the buccalin-positive population.

Among neuropeptides surveyed, only buccalin produced detectable signal in the INC, while FLRIamide, bradykinin-like peptide, and FCAP were all abundant in the ANC but absent from the INC, further supporting a distinct functional role for the INCs. FLRIamide belongs to the FMRFamide-related peptide (FaRP) family, a major peptidergic system across molluscs^47^, and its presence in the ANC but not the INC further differentiates the modulatory repertoires of these two cords. Buccalin-positive cells in the INC did not show detectable vGlut signal, indicating that the peptidergic population is molecularly distinct from the glutamatergic one rather than representing a co-transmitting subpopulation. Buccalin was originally characterized in the Aplysia californica feeding neuromuscular system, where it functions as a modulatory cotransmitter in a cholinergic motor neuron^48–50^. Its expression in the octopus INC appears to follow a different logic, with buccalin-positive cells forming a separate peptidergic class rather than being embedded within a glutamatergic or cholinergic one. The INCs sit directly within the arm musculature, in close proximity to the distinctive neuromuscular junctions of the octopus arm^51,52^, placing a peptide with documented effects on molluscan muscle in a position to modulate local muscle fibers or nearby motor circuits as a standalone peptide signal. Whether this buccalin-positive population projects to different targets than the vGlut-positive population will require higher-resolution spatial analysis and functional work to resolve.

Our finding that a ChAT transcript was not detectable in the INC contrasts with immunohistochemical work ^23^ reporting pChAT-positive structures in the INC of *Octopus vulgaris*. Several non-mutually-exclusive possibilities could account for the difference. The two studies use different species and different molecular targets: Sakaue and colleagues^23^ localized two distinct ChAT proteins, common-type (cChAT) and peripheral-type (pChAT), and reported markedly different distributions for each in the arm. Our HCR probe was designed against a transcript sequence identified in our prior *O. bocki* arm work^14^, and although in principle HCR can detect transcripts from multiple paralogs depending on probe design, we cannot rule out reduced affinity for one paralog over another. It is also possible that pChAT protein persists in INC fibers whose somata and transcripts lie outside the sampled tissue, since immunohistochemistry detects protein along axonal projections rather than at the site of synthesis. Resolving this discrepancy will require targeted probe sets against each ChAT paralog and parallel protein-level validation in *O. bocki*.

### Glial abundance and structural implications

Glial cells are a major constituent of INC tissue, as judged both by HCR marker abundance and by the prevalence of putative glial morphologies in the EM data. The broad spatial distribution of glial markers, extending through the tract rather than being confined to cell body regions, suggests structural and possibly metabolic support of axon bundles and synaptic contacts throughout the cord. This is consistent with general glial roles described in invertebrate nervous systems, including ensheathment, homeostatic support, and interactions with synaptic regions^53–55^, although the specific glia-like cell types in cephalopod peripheral tissues remain incompletely described. The morphologies we describe here, with large flattened cell bodies and branching processes wrapping the cord margins, resemble the putative protoplasmic glia identified in the axial nerve cord of the same species^14^, suggesting that protoplasmic-like glia may be a consistent feature of peripheral arm nervous tissues in octopus, distributed across both central (ANC) and embedded (INC) cords. The morphological diversity of putative glial cells observed in the EM data raises the possibility that the INC contains functionally distinct glial subtypes. Resolving glial identity in this system will require additional molecular markers and, ideally, validation at the protein or functional level.

### The oblique connective is an adjacent but distinct peripheral pathway

A key finding of this study is the structural independence of the OC from the adjacent INC in the sampled high-resolution volume. The oblique connectives were described originally as tracts linking INCs on the same side of the arm at regular intervals, implying that they carry fibers connecting the oral and aboral cords directly. More recent work has shown that OCs course through the arm musculature and extend beyond the INC, potentially crossing the midline of the arm across the oral and aboral aspects. This spiraling organization suggests that rather than being a simple tract connection, they form an independent component of multiple peripheral pathways that coexist in close anatomical proximity^13^. Although their continuation beyond the INCs had been described previously^13^, the close positional relationship between the OC and INC naturally raises the possibility of structural integration. Our data do not support direct integration between these structures in the sampled region. Across the observed OC-INC boundary, a continuous membranous separation was present and no fibers or processes were shared. The OC therefore appears to be an adjacent but distinct peripheral pathway, not a lateral branch or structural extension of the INC.

This finding may have implications for anatomical models of peripheral arm coordination: if the OC and INC do not exchange processes locally, then any coordination mediated by these pathways must involve indirect synaptic relationships or modulatory interactions rather than direct structural continuity at the sampled boundary. The presence of cell bodies, tract fibers, and vesicular boutons within the OC suggests that the OC has cellular and computational complexity of its own, rather than functioning solely as a bundle of connecting fibers.

The absence of detected OC-INC synaptic contacts does not rule out functional interactions at a distance, including diffuse release of neuropeptides or other modulatory substances from one structure acting on receptors in the other. Such modulatory mechanisms are well documented in invertebrate nervous systems^41,56,57^ and cannot be excluded based on structural data alone. However, the absence of direct structural continuity in the region of close contact is informative and should shift future models of INC-OC relationships toward indirect, projection-based, or modulatory mechanisms rather than assuming local tracts merge or that fibers project from one INC, through the OC and into the other INC on the same side.

## Conclusions

The INCs of the octopus arm are more complex and more molecularly diverse than their historical treatment as small peripheral, longitudinal cords might suggest. They contain organized internal compartments, multiple neuronal and glial cell types, and a molecular signature distinct from that of the ANC, indicating that they are not merely structural extensions of central arm pathways but circuits with their own cellular composition. The physical separation of the OC from the INC, despite their close positional relationship, clarifies that these are adjacent but distinct peripheral pathways and shifts questions of peripheral coordination toward understanding OC circuitry in its own right. Together, these findings provide a cellular and molecular foundation for future mechanistic and comparative studies of distributed sensorimotor control in cephalopod arms.

## Acknowledgments

We thank members of the Crook Lab for contributions to animal care, experimental procedures, and scientific discussions. The Anastassov Lab at SFSU provided valuable assistance in preparing tissues for EM and additional expertise in electron microscopy and connectomics, and provided equipment access. We thank Annette Chan and the CMIC at SFSU for assistance with confocal imaging. We thank Grahame Kidd and Emily Benson at the Cleveland Clinic for EM imaging and data management, and the webKnossos team for assistance with alignment, segmentation, data hosting and annotation tools. Confocal imaging was performed on a Leica Stellaris 5 microscope on a DMi8 supported by NSF MRI 2018239. Amy Weir provided illustrations used in Figure 1. This work was supported by a Paul G. Allen Frontiers Group Distinguished Investigator Award in Neural Circuit Design, and NSF CAREER 2047331, to RJC.

