## Supplemental Figure S1 for "Molecular and ultrastructural characterization of the intramuscular nerve cords of the octopus arm"

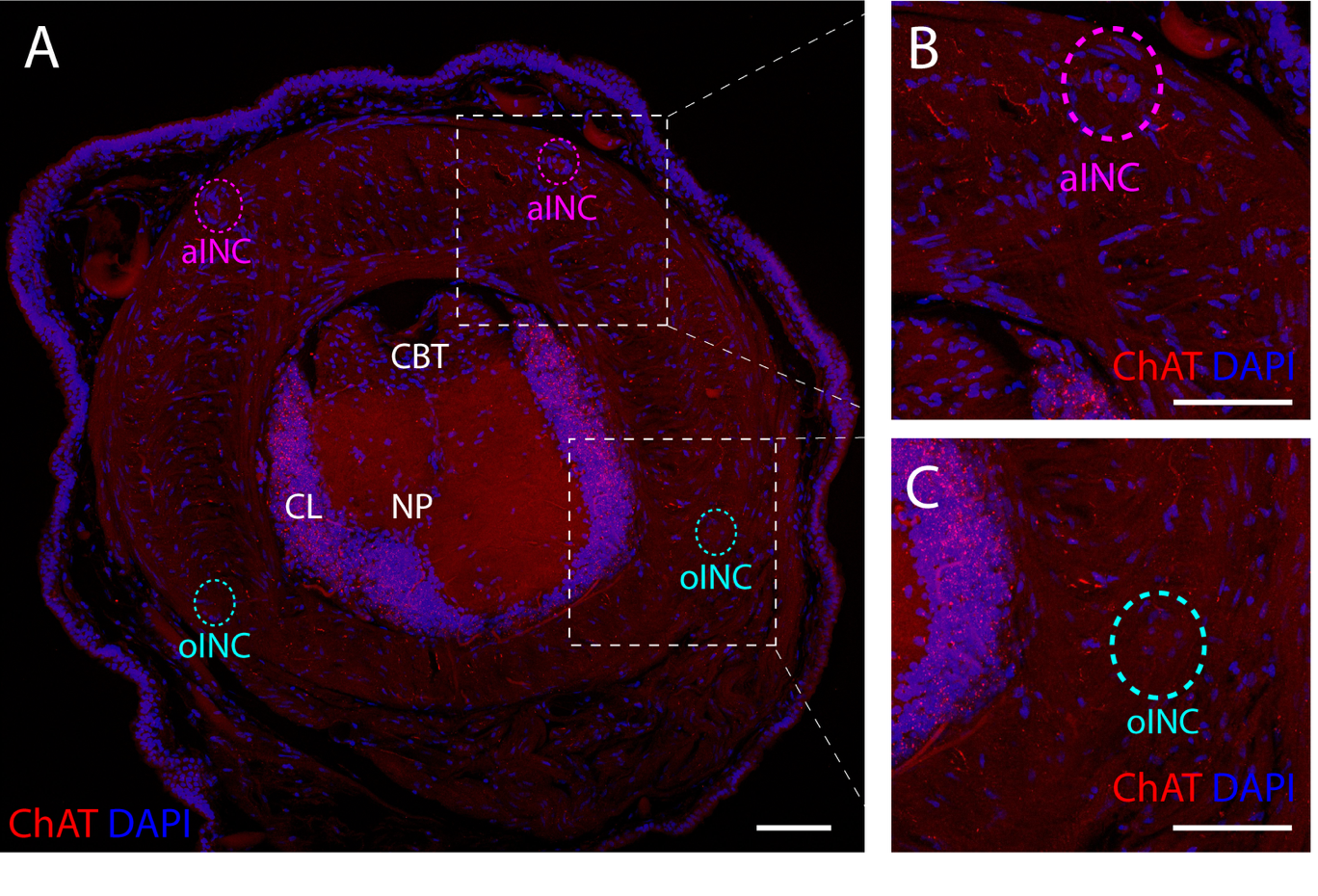

**Figure S1: ChAT is abundant in the axial nerve cord but not detected in the INCs.** HCR labeling for choline acetyltransferase (ChAT, red) with DAPI (blue) in an *Octopus bocki* arm cross section. A. ChAT signal is abundant throughout the axial nerve cord (ANC) but is absent from the two aboral (aINC, magenta) and two oral (oINC, cyan) intramuscular nerve cords circled in the same section. Dashed boxes indicate the regions magnified in B and C. B. Magnified view of an aINC, showing no detectable ChAT signal. C. Magnified view of an oINC lacking ChAT signal. The strong ANC labeling confirms the probe hybridized, indicating that the absence of signal in the INCs reflects a true lack of detectable ChAT transcript rather than failed hybridization. Scale bars: 50 μm.
